# Advances in culturing of the sea star *Patiria miniata*

**DOI:** 10.1101/2024.12.13.628458

**Authors:** Vanessa Barone, Luisa Coronado, Deka Ismail, Sareen Fiaz, Deirdre C. Lyons

## Abstract

The use of the sea star *Patiria miniata* as a model system has produced groundbreaking advances in a disparate set of biomedical research fields, including embryology, immunology, regeneration, cell biology and evolution of development. Nonetheless, the life cycle of *P. miniata* has not yet been closed in the laboratory, precluding the generation of stable transgenic and mutant lines, which would greatly expand the toolset for experimentation with this model system. Rearing *P. miniata* in the laboratory has been challenging due to limited knowledge about metamorphosis cues, feeding habits of juveniles and their relatively long generation time. Here we report protocols to rear *P. miniata* embryos through sexual maturity in a laboratory setting. We provide detailed staging of early embryonic development at different temperatures, and show that larvae can be raised to competence in as little as 15 days. We find that retinoic acid induces metamorphosis effectively and present methods to rear juveniles on commercially available foods. We show that in a flow-through system, juveniles double in size every 2 months and reach sexual maturity in approximately 2 years. We report the first example of *P. miniata* raised through sexual maturity in a laboratory setting, paving the way for the generation of stable mutant sea star lines.

## Introduction

Numerous seminal discoveries in cell and developmental biology have been made using echinoderms as model systems, shaping our understanding of how new life forms^1^. Although the most widely used echinoderm model is certainly the sea urchin, the sea star *Patiria miniata* has emerged as a powerful system for cell biology^2–6^, biophysics ^7,8^ epithelial morphogenesis^9^, evolution of development ^10–12^, regeneration^13^, to name a few. Therefore, a plethora of methods have been established to alter gene expression and protein function during sea star development. These include injection of mRNA, DNA, morpholinos and CRISPR/Cas9 mutagenesis ^2,14^. Most recently, a plasmid-based toolkit has been developed that allows spatio-temporally controlled CRISPR/Cas9 mutagenesis^15^ and a system for easy delivery of constructs in the oocyte^16^.

All these tools, however, rely on delivery of material in the oocyte, zygote or early embryo, limiting the life stages accessible to experimentation and the types of possible experiments. This is because stable lines of mutant and transgenic sea stars are currently not available. Given that efficient methods for mutagenesis exist, the main challenge to the generation of sea star mutant lines is, indeed, establishing laboratory cultures. Before this study, *P. miniata* had not been raised to sexual maturity in a laboratory setting and, therefore, laboratory sea star lines are not available. Two difficulties of establishing laboratory bred sea stars are their bi-phasic life cycle (schematized in **Fig 1**) and their long generation time ^17^. *P. miniata* sea stars are broadcast spawners, planktotrophic indirect developers^18^. Gametes are released into the ocean, where fertilization occurs, followed by development into a bilateral swimming larva, the bipinnaria^19–22^. The bipinnaria feeds on micro-plankton for several weeks before reaching the competent stage, i.e. brachiolaria, and eventually undergoing metamorphosis ^18,23^. The cues that trigger metamorphosis are currently unclear. Metamorphosis has been induced in the lab by introducing biofilms, coralline algae and mollusc shells ^21,23^, but only a small proportion of larvae underwent metamorphosis in these conditions and over a long period of time ^12,23–25^. This poses practical issues when attempting to raise sea stars in the lab, as large numbers of larvae need to be cultured and monitored for metamorphosis for several weeks.

**Fig 1.**
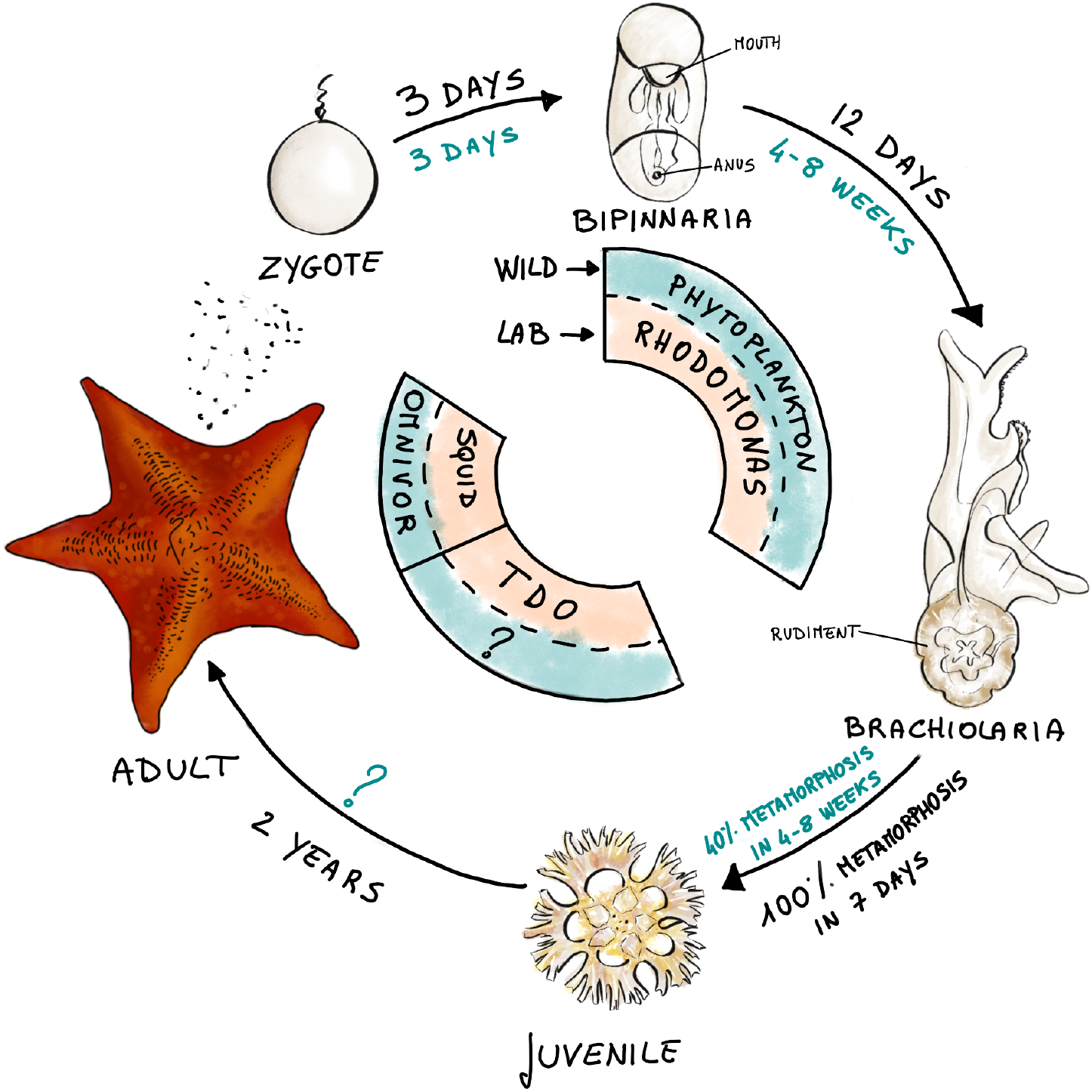
*P. miniata* life cycle. Schematic representation of the main phases of the sea star life cycle. Duration of stages and metamorphosis efficiencies reported in previous studies are highlighted in blue, results from this study in black. The central wheel represents known food sources in the wild and the ones used in this study for laboratory cultures.

With their metamorphosis, the sea stars acquire their adult body plan, with the iconic pentaradial symmetry^12,21^. The juveniles have developed arms, tube feet and feed on surfaces in a manner similar to the adults. However, they are on average 0.6 mm in diameter ^25^, while adults reach sizes of 10-20 cm in diameter ^17^. While juveniles are thought to be herbivores ^26,27^, their diet in the wild is unknown, as observing them in their natural habitats is quite challenging due to their small size. It is also unclear how long it takes for juveniles to reach sexual maturity, i.e. to produce mature gametes, as individual juveniles cannot be tracked over time in the wild. One study on a *P. miniata* population in British Columbia estimated a generation time of two years ^17^. At the time of writing, we have failed to find published reports of *P. miniata* raised to adulthood in captivity.

Here we build on previous work on *P. miniata* ^26^ and other echinoderm species ^26,28,29^ and outline a protocol for larval rearing geared toward speed, low maintenance and efficient metamorphosis. We provide evidence that newly metamorphosed juveniles can survive for long periods of time without feeding but grow only if nutritious substrates are provided. Finally, we show that *P. miniata* can be raised to sexual maturity in a laboratory setting with a generation time of approximately 2 years.

## Results

### Embryonic development at 16°C and 20°C

The early development of *P. miniata* at a temperature of 16°C has been previously described ^19–21,30^. Fertilization is followed by raising of the fertilization envelope and fusion of the female and male pro-nuclei, after which the zygote starts dividing via holoblastic cleavage ^19,20,30^. At these early stages, the first embryonic axis can be identified as the polar bodies are extruded on the side of the embryo that will give rise to anterior ectoderm (animal pole), opposite to the side where the mesoderm and endoderm are specified (vegetal pole)^14,31^. During cleavage, embryonic cells organize into a monolayered epithelium that encircles a cavity, i.e. a blastula^9,14,20,30,31^. Blastulae hatch out of their fertilization envelopes and swim in the water column thanks to their cilia beating^20,30,31^. Shortly after, gastrulation begins, when mesoderm and endoderm progenitors invaginate at the vegetal pole and start forming the gut, also called archenteron^20,30,31^. Invagination continues, the gut keeps protruding inside the blastocoele, while the embryo itself elongates along the antero-posterior axis^20,30,31^. Once the gut has reached approximately the middle of the embryonic cavity, mesenchymal cells ingress from the archenteron and migrate into the cavity^20,30^. The gut then starts bending toward one side of the anterior ectoderm: the mouth opens where the tip of the gut comes into contact with the anterior ectoderm^20,30,31^. At this point the embryo can start feeding on micro-plankton and it is named a bipinnaria larva.

However, detailed staging and comparisons of development and survival rates at different temperatures have not been performed. Development can be accelerated by increasing temperature in many model systems, which can be helpful when aiming at reducing generation times. Therefore, we documented the development of sea star embryos up to the bipinnaria stage at two different temperatures, 16°C and 20°C (**Fig 2**). We split siblings into two groups, placed them in either a 16°C or a 20°C incubator and observed them at regular intervals.

**Fig 2.**
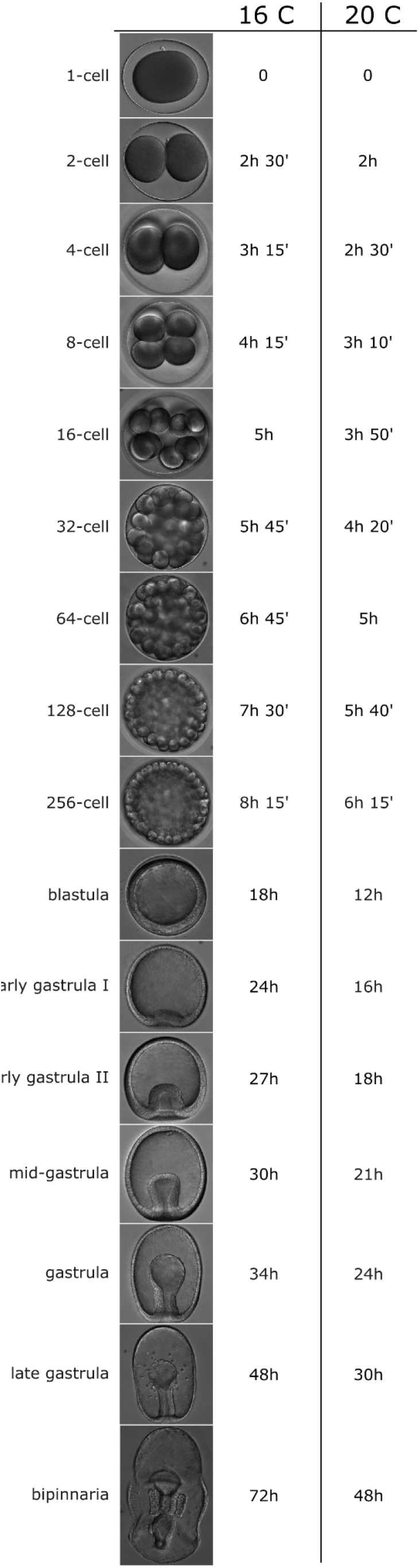
Time-course of *P. miniata* embryo development at 16°C and 20°C. Schedule of development into bipinnaria at 16°C and 20°C.

At 16°C, the first cell division is completed 2.5 hours post fertilization (hpf), and subsequent cleavages occur approximately every 45 min. The embryos are blastulae at 18 hpf, hatch between 20 and 22 hpf and initiate gastrulation at 24 hpf (early gastrula I). At 27 hpf the archenteron has invaginated visibly (∼30% of the gastrula diameter; early gastrula II) and at 30 hpf it extends to the 50% of the gastrula diameter (mid gastrula). At 34 hpf the archenteron has fully extended and the tip has expanded and thinned: the archenteron has now the typical keyhole shape that defines the sea star gastrula. At 48 hpf, mesenchymal cells have ingressed (late gastrula). At 72 hpf the embryos are bipinnariae, as the archenteron has reached the anterior ectoderm, the mouth is open, the anterior coeloms, that will give rise to hydro-vascular system, have extended and the left posterior coelomic pouch is visible. These results are in line with previous reports ^20,30^.

As expected, development proceeds faster at 20°C (**Fig 2**). The first cell division is completed 2 hpf and cleavages occur approximately every 30 min. Embryos are blastulae at 12 hpf, and initiate gastrulation at 16 hpf. Gastrula stage is reached at 24 hpf and mesenchymal cells have ingressed by 30 hpf. Larvae are bipinnariae by 48 hpf, one full day earlier than their siblings raised at 16°C. These results provide a detailed schedule of key developmental stages at two temperatures.

### Efficient larval culture and induction of metamorphosis

Having established a developmental schedule and survival rates for early embryos, we established protocols for effective rearing of *P. miniata* larvae through metamorphosis. Once *P. miniata* embryos have opened a mouth they begin feeding on microorganisms in the sea water^23,24^. With feeding, the bipinnaria larvae continue developing until they are competent to metamorphose, i.e. they have reached the brachiolaria stage, characterized by the growth of larval arms, and of the image of the future juvenile, the rudiment, on the posterior left side of the brachiolaria ^12,30^. The rudiment is the first instance in which the characteristic pentaradial symmetry of the sea star can be observed, as the juvenile arms and mouth can be discerned in the rudiment ^30^. Once the rudiment is developed, the larvae are ready to undergo metamorphosis: they adhere to an appropriate substrate with the anterior arms, the larval body is reabsorbed and transformed into the juvenile ^12,21,23–25,30^.

Previous work identified *rhodomonas* unicellular algae as an efficient food source for *P. miniata* larvae ^24,26^, and larval cultures have been raised to competency at 16°C in 4-8 weeks ^21,23,24,26^. We modified previous protocols for larval rearing as follows by i) raising larvae at 20°C, ii) feeding higher amounts of *rhodomonas sp*., iii) culturing the larvae in filtered sea water (FSW) with antibiotics and iv) reducing the frequency of water changes (see Methods).

We found that larvae developed to the late bipinnaria stage by 9 days post fertilization (dpf) (**Fig 3A**) and to the brachiolaria stage by 14 dpf, when the rudiment became visible (**Fig 3A**). We observed no obvious further development between 14 and 21 dpf (**Fig 3A**). We then measured larval length as a proxy of growth and found that larvae reached their maximum size (1.54 ± 0.13 mm) by 14 dpf (**Fig 3B**). Survival rates for larvae cultured in these conditions were consistently above 90% at 14 dpf and above 80% at 21 dpf (**Fig 3C**).

**Fig 3.**
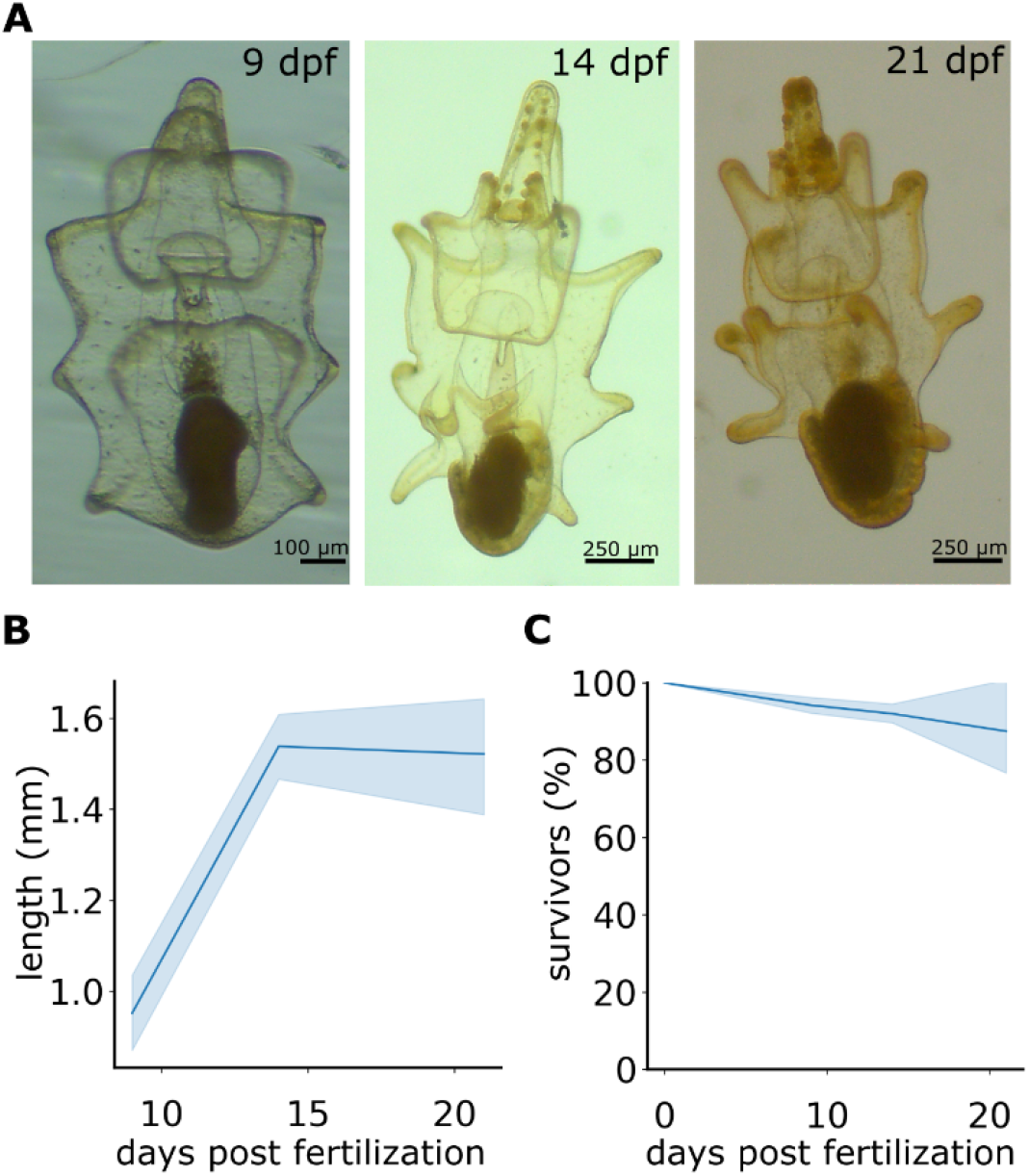
Larval growth on excess food. **A**) Representative images of *P. miniata* larvae fed on Rhodomonas at a concentration of 10’000 cells/ml for 9, 14 and 21 days. Fertilized embryos were raised in petri dishes PS-FSW at 16°C for 3 days, when they reached the bipinnaria stage. They were then transferred to beakers with gentle agitation, incubated at 20°C and fed daily with rhodomonas to a final concentration of 10’000 cells/ml. Larval length **B**) survival (**C**) were quantified. 640 larvae, 4 experiments. dpf: days post fertilization.

We then sought to establish a method to induce reliable and efficient metamorphosis of competent larvae. Given that retinoic acid (RA) has been shown to induce metamorphosis in *Patiria pectinifera*^*32*^, we tested its efficacy on *P. miniata* larvae. To this aim we raised *P. miniata* larvae to 15 or 21 dpf and then exposed them to either DMSO (control) or 0.1 μM RA for 48h. The larvae were then transferred to FSW and metamorphosis was assessed 7 days after first exposure to DMSO or RA. While virtually no control larva metamorphosed, over 80% of 15 dpf and 100% of 21 dpf larvae underwent metamorphosis (**Fig 4A,B**).

**Fig 4.**
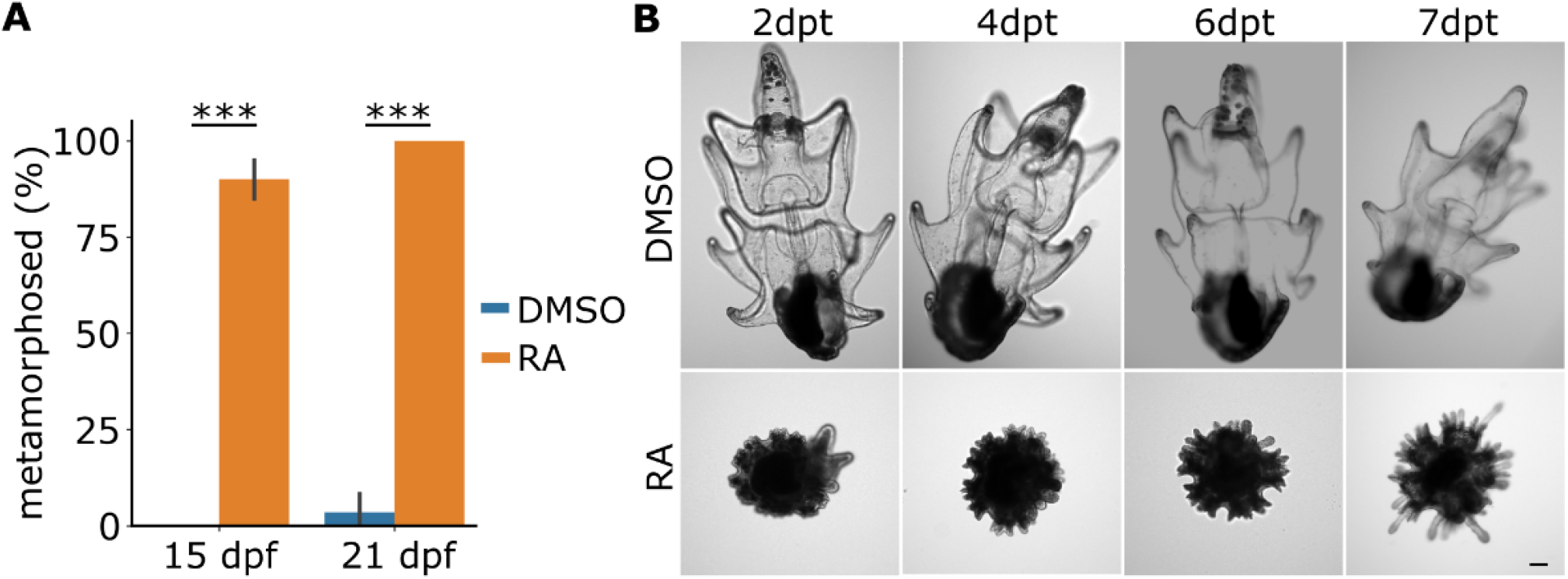
RA induces metamorphosis in *P. miniata* larvae. **A**) Representative images of larvae treated with either DMSO or 0.1 μM retinoic acid (RA). Larvae were cultured for either 15 or 21 days, transferred to 6-well plates with 5-10 larvae each and treated with either DMSO or RA for 48h. Larvae were then transferred to PS-FSW and cultured for 5 days. At 7 days post treatment, the proportion of larvae that underwent metamorphosis was quantified. (**B**). DMSO larvae were not fed during the treatment. 230 larvae, 4 experiments, 3 technical replicates for experiment. Statistical test: Two-way anova, Bonferroni multiple comparison correction. ***: p-value<0.001. dpt: days post treatment. Scale bar 100 μm.

Taken together, these results show that *P. miniata* larvae can be raised at 20°C, they reach competency in 15 days and metamorphosis can be effectively induced by exposure to RA.

### Food sources for *P. miniata* juveniles

The juvenile sea star has the adult body plan but no developed gonads^12,17^. It will feed and grow considerably before becoming a sexually mature adult^17^. One challenge to lab rearing of *P. miniata* is identifying an appropriate food source for newly metamorphosed juveniles, as the small size of the juveniles makes observing their behavior in the wild difficult. When juveniles start feeding and on what they feed is currently unknown. Therefore, we tested survival and growth of juveniles provided with food sources that could potentially be used for laboratory culturing of *P. miniata*. To this end, we split newly metamorphosed juveniles (1 week post metamorphosis, wpm) into 4 groups and transferred each group to a 10 mm petri dish containing FSW and either i) no food (CTRL), ii) rhodomonas sp., iii) a glass slide that had been incubated in local aquaria for at least 3 weeks (biofilm) or iv) several fragments of crustose coralline algae (CCA). We then cultured the juveniles in the petri dishes for up to 4 months (**Fig 5A**). We performed weekly water changes, as well as measurement of survival and growth (**Fig 5B,C**). Growth was estimated by measuring the diameter of 5-10 juveniles per dish.

**Fig 5.**
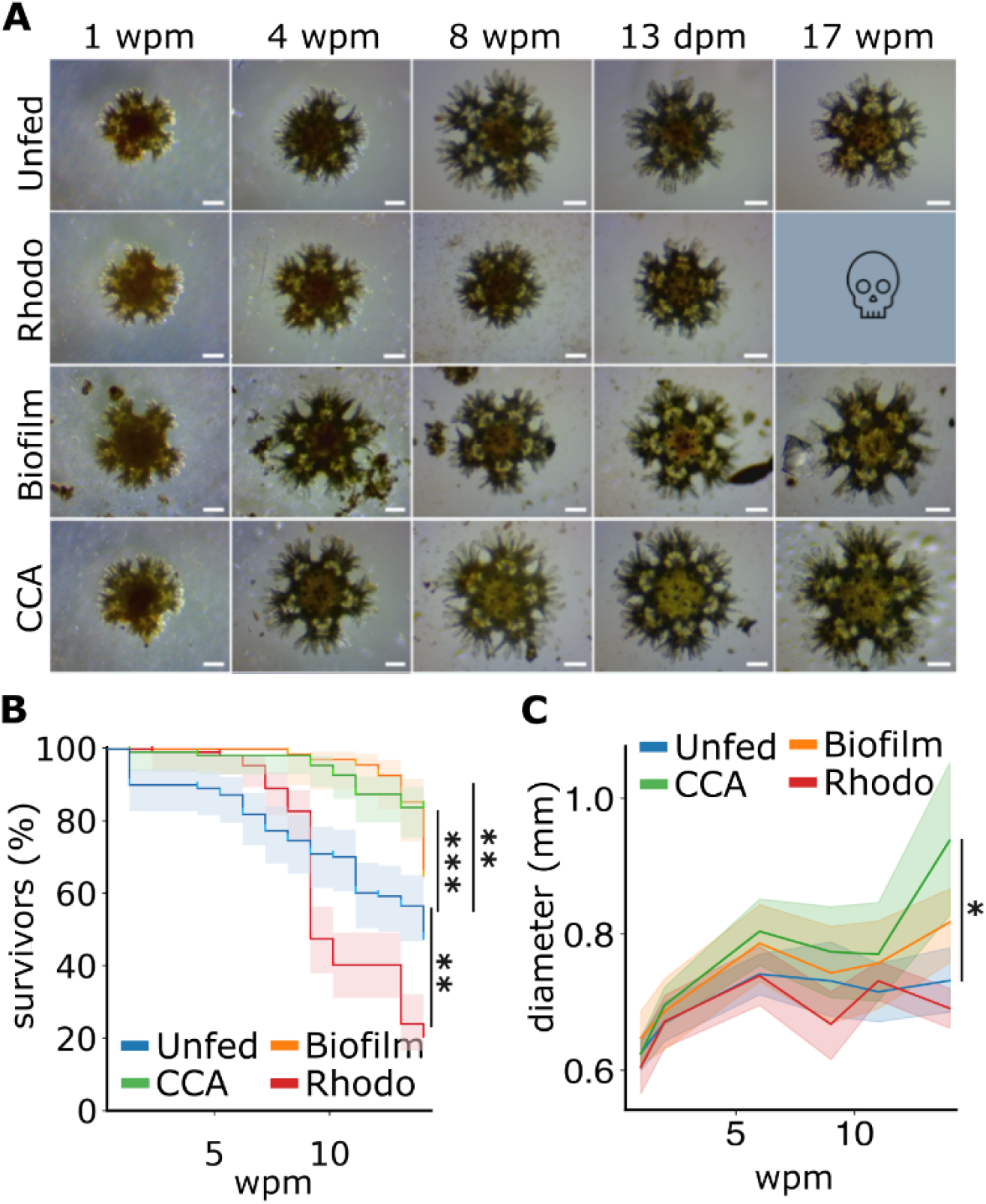
CCA sustains minimal growth of *P. miniata* juveniles. **A**) Representative images of juveniles cultured in petri dishes and provided with no food (unfed), rhodomonas (Rhodo), a glass slide covered with natural biofilm (biofilm) and crustose coralline algae (CCA). Survival (**B**) and growth (**C**) were quantified. wpm: weeks post metamorphosis. 441 juveniles, 4 experiments. Error bars: 95% CI. Statistical test on survival: Mantel-Cox test with Bonferroni multiple comparison correction. Statistical test on growth: one-way anova on last measured timepoint, Tukey pairwise comparison and Bonferroni multiple comparison correction; *: p-value <0.05; **: p-value <0.01; ***: p-value <0.001 Scale bars: 50 μm.

We found that juveniles can survive without any food source for several weeks, with survival rates in control conditions declining sharply only after 6 weeks (**Fig 5B**). The addition of rhodomonas has a negative effect on both survival and growth (**Fig 5B,C**). Juveniles fed with biofilm showed improved survival compared to controls, but similar growth (**Fig 5B,C**). The addition of CCA, instead, improved both survival and growth (**Fig 5B,C**). Interestingly, growth of juveniles fed with CCA was faster than controls but seemed to plateau after 4 weeks (**Fig 5C**). We reasoned that grazing on CCA may sustain the first phases of juvenile growth but is insufficient for further development. Therefore, on week 10 we added a small piece of squid to the CCA cultures, which was removed after 24h to avoid excessive fouling. The size of CCA juveniles increased slightly over the following 5 weeks (**Fig 5C**).

These results show that juveniles can survive without substantial food for long periods of time and that they can start feeding shortly after metamorphosis. However, the growth rates in these experiments were somewhat disappointing, as in the best condition juveniles only doubled in size after 4 months. This is probably due to the lack of key nutrients.

We then turned to testing commercially available foods for aquaria and mariculture, but found that fouling precluded testing of these food sources in the petri-dish cultures. However, we did identify one product (TDO CHROMA BOOST™, Reef Nutrition) which the juveniles grazed on (**Fig 6B**), therefore we used TDO to feed juvenile cultured in a flow-through system (**Fig 6**).

**Fig 6.**
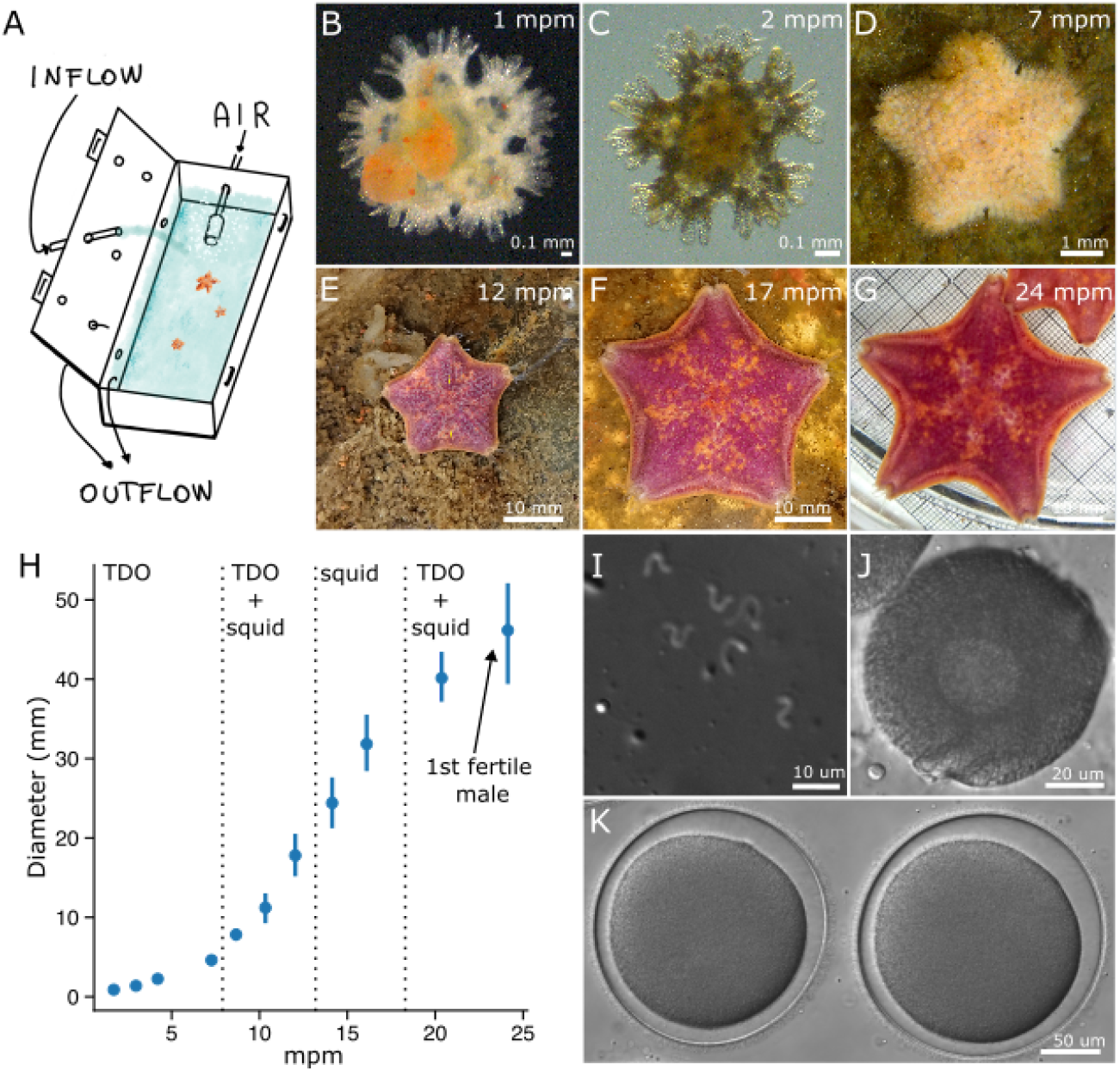
Culture of *P. miniata* juveniles to sexual maturity. A) Schematic representation of the flow-through system used for culturing juveniles. B-G) Representative images of growing juveniles. B) shows a ventral view of a juvenile feeding on TDO. H) Quantification of juvenile growth: 5-10 juveniles fed on TDO were photographed and their diameter measured to monitor growth. I) DIC image of sperm from the first fertile male. J) DIC image of an immature oocyte isolated from one juvenile at 24 months post metamorphosis. K) Zygotes generated by fertilizing mature oocytes with sperm isolated from the first lab cultured fertile male of *P. miniata*. One experiment started with 100 juveniles at one week post metamorphosis. mpm: months post metamorphosis.

### Culture of juvenile *P. miniata* to sexual maturity

A flow-through culturing system, with far greater and adjustable water exchange rates, allows to provide larger amounts of food to cultures, which may be pivotal to raise *P. miniata* juveniles to adulthood. The main challenge to implement such a flow-through system is the small size (0.6 mm in diameter, **Fig 5**) and flexibility of the newly metamorphosed juveniles: sea star juveniles, just as sea star adults, can squeeze through openings much smaller than their diameter. The challenge is then to have meshes small enough to keep the juveniles from escaping, while also allowing sufficient water flow. To test the feasibility of such an approach, we devised a low cost system using perforated scrapbook boxes, equipped with meshes on the sides (**Fig 6A**). Each box was equipped with a sea water inflow and an air bubbler (**Fig 6A**).

We placed 100 newly metamorphosed juveniles (1 wpm) in each of 3 boxes. One box was filled with a bed of CCA, one with *Ulva lactuca* and one was left empty. In addition, juveniles were fed weekly with TDO pellets. For several months, we were not able to find juveniles in the boxes with substratum (CCA and Ulva), so we did not monitor their growth or survival. At semi-regular intervals, 5-10 juveniles were taken from the box with no additional substratum and photographed to monitor their growth (**Fig 6C-G**). We found that juvenile size doubled approximately every 2 months (Fig 5H). At 7 months post metamorphosis (mpm), we added squid to the juveniles diet (**Fig 6H**). At 14 mpm, we removed the TDO from the diet and fed the growing juveniles solely with squid (**Fig 6H**). While growth continued, it slowed down (**Fig 6H**) and some of the juveniles started to lose pigment. Therefore, at 18 mpm we re-introduced the TDO to the diet (**Fig 6H**). At 18, 22 and 24 mpm we dissected the surviving juveniles (8) to test for gonad development. At 24 mpm we found one fertile male (**Fig 6I**) and two females with immature oocytes (**Fig 6J**). The sperm of the fertile male successfully fertilized mature oocytes dissected from adult females (**Fig 6K**). At 24 mpm when the experiment was terminated, 8 of the 100 juveniles in the box survived, one had reached sexual maturity and 2 had gonads but immature gametes. These results establish that *P. miniata* can be reared in captivity.

## Discussion

The advent of CRISPR/Cas9 has opened the doors for directed mutagenesis in a wide range of organisms that can now become fully genetically enabled. The generation of mutant and transgenic lines for echinoderms is now a reality and is already expanding the realm of mechanistic experimentation that is now possible in the sea urchin^28,33–37^.

To fully exploit the power of the echinoderm systems, however, it will be important to have at least one model species for each of the echinoderm classes, namely echinoids, asteroids, holothuroids, ophiuroids and crinoids, for which mutant and transgenic lines are available. Once that goal is achieved, we will be able to fully explore the biological diversity found in this heterogenous group of marine animals.

We present evidence for the first time that *P. miniata* can be reared to adulthood in captivity, and provide a timetable of embryonic development at 16°C and 20°C, protocols for efficient larval culturing, and metamorphosis induction. After the bipinnaria stage, we cultured larvae at 20°C with very high survival rates (above 80%), which allowed the larvae to reach competency considerably faster than previously reported (2-3 weeks compared to 4-8 weeks^21,23–25^). To maximize survival and minimize culturing times, our suggestion is that embryos be cultured at 16°C until 72hpf and then transferred to 20°C until they reach competency.

We identified exposure to commercially available retinoic acid (RA) as an efficient method to induce metamorphosis, as we found that 100% of competent larvae simultaneously metamorphosed after incubation with 0.1μM RA. This is a higher percentage of metamorphosis than previously reported for metamorphosis induced by exposure to natural substrates ^17,21,23–25^, which resulted in at best 40% of larvae metamorphosing ^23^, and in sibling larvae initiating metamorphosis at different times over a period of 4-8 weeks ^24,25^. For culturing purposes, the RA method has the advantage of producing batches of simultaneously metamorphosed juveniles that can be transferred to aquarium systems for juvenile rearing. However, it is important to note that RA-induced metamorphosis differs substantially from naturally occurring metamorphosis. For instance, *P. miniata* larvae have been shown to metamorphose upon adhesion to substrates (such as pebbles or CCA covered with natural biofilms) ^23^, while RA induced larvae do not adhere to the substrate and continue moving via ciliary beating until metamorphosis is completed. Naturally-induced metamorphosis is also faster than RA-induced metamorphosis ^21^. RA has been shown to induce metamorphosis in *P. pectinifera* sea stars ^32^ and it is thought to be an effector of the metamorphosis developmental program, probably triggered by environmental cues ^32^. As of now, the nature of these environmental cues is unclear for *P. miniata*. While efforts are underway to identify a natural compound that induces metamorphosis consistently and efficiently, we think that the use of RA is a useful method to streamline lab culturing efforts at this time. Inducing metamorphosis with RA has allowed us to experiment with culturing of *P*.*miniata* juveniles, which had been previously more challenging due to the unpredictability of metamorphosis rates.

We show here that *P. miniata* juveniles can be easily maintained in the lab by housing them in petri dishes for several months. We performed weekly water changes, as pilot experiments performing water changes daily or every two days resulted in low survival (data not shown). We noticed that water changes often result in juveniles detaching from the substrate, and hypothesize that it may be a source of stress to the animals. We also show that juveniles can survive for extended periods of time without any substantial food source, as survival in unfed controls started declining only after four weeks, and 49.72% ± 36.24% juveniles had survived after 14 weeks, when the experiment was terminated. Survival was hampered by the weekly addition of *Rhodomonas sp*. to the petri dishes: it is possible that the high concentration of algae combined with infrequent water changes created anoxic conditions. Presence of a biofilm, instead improved survival, which started to decline only after 12 weeks in culture. The biofilm, however, was not sufficient to support growth. Presence of crustose coralline algae (CCA) in the dishes resulted in survival rates above 90% for the entirety of the experiment and in modest but measurable growth. Juveniles fed with CCA showed growth in the first four weeks, followed by a plateau for about 5 weeks. In week 10, we added a small piece of squid to the CCA petri dishes and observed further growth in the following weeks. Therefore, it is probable that juveniles are capable of feeding immediately after metamorphosis and that their survival and speed of growth greatly depends on the type of food available. In this experiment we could not easily observe clear grazing stripes, and we could not distinguish whether the juveniles were feeding on the CCA itself or on biofilms growing due to the presence of CCA. Either way, additional food sources are needed for further growth. This setup however, could be useful for researchers interested in maintaining young juveniles in the lab for experiments that do not necessitate fast growth, such as behavioral or toxicological assays.

To achieve further growth, we turned to artificial foods and a flow-through culturing system: this allowed us to raise juveniles to sexual maturity. This is an exciting result, as it demonstrates that *P. miniata* can be reared in the lab, paving the way for establishing laboratory strains for this model system. However, the first male to reach sexual maturity did so after 2 years and overall survival was very low (8% for this experiment). There is much room for improvement.

For example, in this experiment, we used minimally filtered sea water at 16°C. This resulted in a layer of biofilm and detritus accumulating in the tanks, which was challenging to clean out without removing juveniles attached to the bottom: this may have reduced survival. Our tank design was also suboptimal, as we noticed juveniles were able to escape the scrap-boxes until they reached a diameter of about 8 mm. Therefore, low survival rates may also be due to loss of escapist juveniles. We also made the choice to remove the TDO from the juvenile’s diet once we switched to feeding them with squid, which may have slowed down growth. Finally, it is possible that raising the juveniles at higher temperatures, up to 20°C, would considerably reduce the time needed to reach sexual maturity. Improving on the methods presented here will permit the engineering of transgenic lines of *P. miniata*, which are sorely needed to push the boundaries of the research supported by echinoderm model systems.

## Methods

### Adult animal husbandry and embryo culture

Adult *Patiria miniata* were purchased from Monterey Abalone Company (Monterey, CA) or South Coast Bio-Marine LLC (San Pedro, CA) and held in free flowing seawater aquaria at a temperature of 12-16°C. Sea star gametes were obtained as previously described^26^. Briefly, ovaries and spermogonia were dissected via a small incision on the ventral side of adults. Sperm was stored undiluted at 4C while ovaries were fragmented to release oocytes in FSW. Maturation of released oocytes was induced by incubating for 1h at 16°C in 3 μM 1-Methyladenine (Fisher Scientific, 5142-22-3). All embryos were raised in 0.22 μm filtered sea water (FSW) with the addition of 0.6 μg/ml Penicillin G sodium salt (Millipore Sigma, P3032) and 2 μg/ml Streptomycin sulfate salt (Millipore Sigma, S1277).

### Embryo survival experiments

For each mating pair, the same number of zygotes (50-100) were transferred to two 60 mm petri dishes containing PS-FSW. One petri dish was placed in a 16°C incubator and the other in a 20°C incubator. At 48hpf, the 20°C petri dish was examined and surviving larvae were counted. At 72hpf, the 16°C petri dish was examined and surviving larvae were counted. Embryos that did not open a mouth, or showed obvious morphological defects were not considered survivors.

### Larval cultures

Healthy early bipinnaria larvae were selected at 72 hpf (16°C) and transferred into 500 ml beakers filled with PS-FSW at a concentration of 0.3 larvae/ml.

Larvae were kept in gentle agitation using a paddle connected to a slow rotor (5-10 rpm), as previously described [ref] and fed rhodomonas daily at a final concentration of 10000 cells/ml. A complete water change was performed weekly, by picking and transferring all surviving larvae into a clean beaker. Concomitantly, larvae were counted to assess survival and a subset was photographed and measured to assess growth. Growth was measured as follows. Three larvae per beaker were transferred in double strength seawater for 30 sec and then back to normal FSW. This procedure damages cilia and prevents the larvae from moving for about 20 min. Larvae were then photographed using a Zeiss Semi 305 stereoscope. Larval growth was measured using Fiji (ref) by recording the length of the larva from the tip of the anterior arm to the posterior end.

### Metamorphosis

Metamorphosis of competent larvae (15 or 21 dpf) was induced with RA. To assess metamorphosis efficiency, 10 larvae were transferred to each well of a 6-well plate containing 10 ml of FSW with the addition of either DMSO (CTRL) or 0.1 uM RA. Larvae were incubated for 48h and then the drug was washed out. Metamorphosed juveniles were counted once ciliary movement ceased, i.e. 7 days post treatment. Juveniles were classified as metamorphosed if they had developed tube feet and adhered to the substrate.

### Juvenile cultures: focal cultures

Metamorphosed juveniles (7 days post treatment) were transferred to 10 mm petri dishes containing FSW (without antibiotics). Juveniles originating from the same mating pairs were split into 4 petri dishes, each containing 30-50 juveniles. A different food source was then added to each petri dish, i.e. none (CTRL), 1 ml of rhodomonas culture (Rhodo), a glass slide that had been placed in one of the adult aquaria for 4-weeks (Biofilm) and a few fragments of coralline crustose algae (CCA). Every week, FSW was exchanged, food refreshed, surviving juveniles counted and 5-10 juveniles per dish were photographed to measure their size.

### Juvenile cultures: flow through system

Metamorphosed juveniles (7 days post treatment) were transferred to 10 mm petri dishes containing FSW (without antibiotics). The petri dish was then placed inside a scrap-box that had been previously prepared. Food safe scrap-boxes were purchased on Amazon and perforated using a power drill, making holes of 0.6 cm in diameter, six on the lid, two on one of the long sides and one hole on one of the short sides of each box. The holes in the long side were covered with a 400 um nylon mesh. The hole on the short side was used to insert a bubbler and one of the holes on the lid was used to insert a sea water line. The scrapbooks were placed on a sea table and sea water filtered to 20 μm was let flow through them for 3 weeks prior to juvenile transfer. This created a biofilm in the scrap-boxes and quickly clogged the nylon meshes. Water was then flowing out of the boxes via the open holes in the lid. 100 newly metamorphosed juveniles (1 wpm) were placed in each of 3 boxes. One box was filled with a bed of CCA, one with ulva algae and one was left empty. In addition, juveniles were fed weekly with TDO CHROMA BOOST™ X-SMALL (Reef Nutrition) pellets for 14 months. After that, juveniles were fed with pieces of frozen squid until 18 months post metamorphosis, when the TDO was reintroduced in the diet. For several months, we were not able to find juveniles in the boxes with substratum (CCA and Ulva), so we did not monitor their growth or survival. Approximately every four weeks, 5-10 juveniles were taken from the box with no additional substratum and photographed to monitor their growth.

### Larval and juvenile size measurements

Larvae were transferred in double strength seawater for 30 sec and then back to normal FSW. This procedure damages cilia and prevents the larvae from moving for about 20 min. Larvae were then photographed using a Zeiss Stemi 305 stereoscope. Larval size was measured using Fiji ^38^ by recording the length of the larva from the tip of the anterior arm to the posterior end. Juveniles were photographed using a Zeiss Stemi 305 stereoscope and, at later stages, using a cellphone camera to photograph juveniles placed on top of mm paper. Juvenile size was measured with Fiji, by measuring the area of an ellipse including the juvenile. The diameter was calculated considering the area of the ellipse as if it were the area of a sphere. We preferred this method to measuring the longest axis of the ellipse because moving juveniles can sometimes be considerably stretched in one direction, so that the major axis of a fitted ellipse does not reflect the size of the animal as much as the overall surface area of the ellipse.

### Statistical analysis

Statistical analyses of data were performed using Python or GraphPad scripts as indicated in the figure captions. Two-way anova with Bonferroni multiple comparison correction was used to test the effects of RA on induction of metamorphosis (Fig 3). Mantel-Cox test with Bonferroni multiple comparison correction was used to compare juvenile’s survival on different food sources (Fig. 4). One-way anova on juveniles sizes at the last measured time point, Tukey pairwise comparison and Bonferroni multiple comparison correction was used to compare growth of juveniles on different food sources (Fig 4). No statistical method was used to predetermine sample size, the experiments were not randomized and the investigators were not blinded to allocation during experiments and outcome assessment.

## Acknowledgements

The authors would like to thank the experimental aquarium facilities at Scripps Institution of Oceanography, and the students of the Development and Cell biology lab course 2019 at UCSD for their help with the metamorphosis experiments. We extend our thanks also to Zak Swartz for sharing information about commercially available foods to be tested, and to all members of the Lyons and Hamdoun labs for sharing their knowledge and experience on culturing echinoderms.

## Authors’ contributions

VB conceived the projects, performed experiments, wrote the manuscript and secured funding. DI, LC and SF performed experiments. DCL secured funding, organized and led the developmental and cell biology course at UCSD -where the students performed experiments included in this manuscript - provided feedback on the project and edited the manuscript.

## Availability of data and materials

All data is included in the manuscript and all materials are commercially available.

